# The transcribed intergenic regions exhibit lower frequency of nucleotide polymorphism than the untranscribed intergenic regions in the genomes of *Escherichia coli and Salmonella enterica*

**DOI:** 10.1101/2022.08.07.503086

**Authors:** Pratyush Kumar Beura, Piyali Sen, Ruksana Aziz, Siddhartha Shankar Satapathy, Suvendra Kumar Ray

## Abstract

The temporary exposure of single-stranded regions in the genome during the process of replication and transcription makes the region vulnerable to cytosine deamination resulting higher rate of C→T transitions. Intra-operon intergenic regions undergo transcription along with adjacent co-transcribed genes in an operon, whereas inter-operon intergenic regions only undergo replication. Hence these two types of intergenic regions (IGRs) can be compared to find out the contribution of replication-associated mutations (RAM) and transcription-associated mutations (TrAM) towards bringing variation in genomes. In our work, we performed a polymorphism spectra comparison between intra-operon IGRs and inter-operon IGRs in genomes of two well-known closely related bacteria such as *Escherichia coli* and *Salmonella enterica*. In general, the size of intra-operon IGRs was smaller than that of inter-operon IGRs in these bacteria. Interestingly, the polymorphism frequency at intra-operon IGRs was 2.5-fold lesser than that in the inter-operon IGRs in *E. coli* genome. Similarly, the polymorphism frequency at intra-operon IGRs was 2.8-fold lesser than that in the inter-operon IGRs in *S. enterica* genome. Therefore, the intra-operon IGRs were often observed to be more conserved. In the case of inter-operon IGRs, the T→C transition frequency was a minimum of two times more than T→A transversion frequency whereas in the case of intra-operon IGRs, T→C transition frequency was similar to that of T→A transversion frequency. The polymorphism was purine biased and keto biased more in intra-operon IGRs than the inter-operon IGRs. In *E. coli*, the Ti/Tv ratio was observed as 1.639 and 1.338 in inter-operon and in intra-operon IGRs, respectively. In *S. enterica*, the Ti/Tv ratio was observed as 2.134 and 2.780 in inter-operon and in intra-operon IGRs, respectively. The observation in this study indicates that transcribed IGRs might not always have higher polymorphism frequency than the untranscribed IGRs. The lower polymorphism frequency at intra-operon IGRs might be attributed to different events such as the transcription-coupled DNA repair, sequences facilitating translation initiation and avoidance of rho-dependent transcription termination.

## Introduction

Many of the functionally related bacterial genes are transcribed as a polycistronic unit. As most of the genes are present under operon units in prokaryotes, two types of untranslated intergenic regions (IGRs) could be found in their genomes; intra-operon IGRs and inter-operon IGRs. Inter-operon IGRs are found in between two separate operons/cistron units. Intra-operon IGRs are found between two adjacent open reading frames in an operon (Figure 1), which is popularly known as Intercistronic regions (ICRs). Inter-operon IGRs are the units that undergo only replication, but intra-operon IGRs are the units that undergo replication as well as transcription, but neither of them codes for protein. The size of inter-operon IGRs is larger than intra-operon IGRs. The small-sized intra-operon IGRs are evolutionarily preferred only to prevent energy wastage in transcribing longer intra-operon IGRs. Sometimes co-transcribed genes can also be found to be overlapped by a few sequences, but they lack intra-operon IGRs. The bacterial chromosome has only 10-15% regions known as untranslated intergenic regions (IGRs) and contains many regulatory elements with key functions (Thorpe et al. 2017; Sridhar et al. 2011). In past years, intra-operon IGRs were found to be playing an important role in the formation of hairpin loops as well as Ribosome binding sites (RBS) in bacteriophages (Romantschuk and Müller, 1983). Intra-operon IGRs were also found to be important in assigning RBS and recruitment of EF-G gene ((Post and Nomura, 1980)

**Figure 1.**
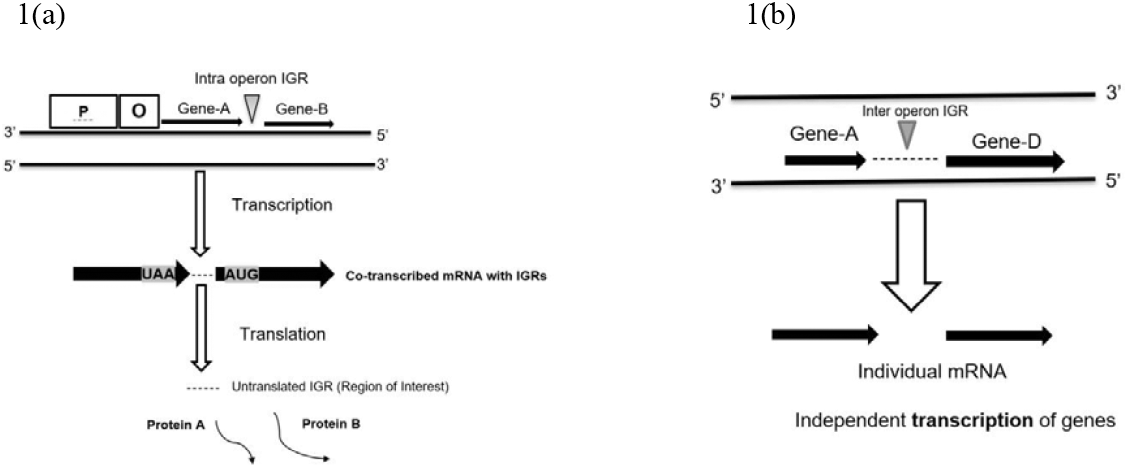
A schematic diagram elucidating the difference in intra-operon IGRs and inter-operon IGRs. Legend. figure 1(a) shows the schematic diagram of an operon P-promoter and O-operator followed by structural regions with genes, between Gene-A and Gene-B the intra-operon IGRs can be found which are transcribed but not translated. Whereas figure 1(b) shows the inter-operon IGRs in between two separate genes/operons, where independent transcription of the operon/genes occur. Usually, the inter-operon IGRs are not transcribed.

Among different types of polymorphism, base substitution occurs predominantly in a bacterial genome apart from insertion/deletion (INDELs). Base substitution is also studied by scientists to a great extent by using *Escherichia coli* as a model organism in recent decades. The temporary exposure of single-stranded regions in the genome during the events like replication and transcription makes the region vulnerable to base substitutions. Scientists have discovered that cytosine deamination is a major reason behind the base substitution in genomic DNA alongside Guanine oxidation leading to G→A substitution. (Kino and Sugiyama 2001; Rocha et al. 2006;Bhagwat et al. 2016). Similarly, the single-stranded exposure of the non-template strand during transcription makes it prone to C→T base substitutions in the non-template strand has been described recently (Francino and Ochman 1997; Mugal et al. 2009). Hence C→T/G→A is a major contributor to the polymorphism spectra in CDS as well as non-CDS regions in chromosomes. The base substitutions are studied as transition and transversion. As nucleotides are divided into two classes; Purine (R) and Pyrimidine (Y), The intra-class substitution between nucleotides is known as Transition (Ti) (R→R, Y→Y). The inter-class substitution between nucleotides is known as Transversion (Tv) (R→Y, Y→R) (Figure 2). Like coding sequences (CDS), substitutions in IGRs can also affect the expression and regulation of the CDS in pathogenic bacteria (Laabei et al. 2014; Casali et al. 2014)

**Figure 2.**
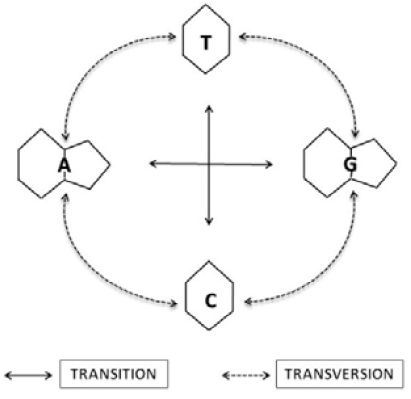
A schematic diagram elucidating transition and transversion in chromosomes. Legend. Purines (A & G) are shown as double-ringed nitrogenous bases and Pyrimidines (T & C) are shown as single-ringed nitrogenous bases. The base substitution between two similar classes of nucleotides is shown as a transition (solid lines with arrows). The base substitution between two different classes of nucleotides is shown as transversion (dotted line with arrow)

As mentioned earlier, intra-operon IGRs undergo transcription which makes them fundamentally different from inter-operon IGRs. We studied a comparative polymorphism analysis using computational tools to find out the different effects of transcription and replication on polymorphism in both these regions. As the frequency of transcription is higher as compared to a standard replication cycle of any bacteria, the regions containing intra-operon IGRs are likely to be exposed as a single strand more frequently than inter-operon IGRs. Hence, we presumed to get a higher polymorphism frequency value in intra-operon IGRs, and to study the possible effect of transcription-induced polymorphism and/or selection in the IGRs of a bacterial genome. In this study, we considered 157 strains of *E. coli* and 366 strains of *S. enterica* and collected 134 and 89 intra operons IGRs respectively for both species, then a comparison with inter-operon IGRs was done for each species.

## Materials and methods

### Finding inter-operon intergenic regions

In this study, we have considered 157 strains of *Escherichia coli* (*E. coli*) and 366 strains of *Salmonella enterica* (*S. enterica*) (Thorpe et al., 2017). We retrieved the inter-operon IGRs from our available dataset referring to the co-ordinate details of the adjacent operons/genes. In our work, we did not consider the possible promoter and terminator regions of the adjacent genes, to avoid any biases in selection. Suppose we found 570 bp inter-operon IGRs, we did not consider 35 bp at either end (70 bp total) for further steps, hence only 500 bp was considered as inter-operon IGRs for that region. This 35 bp at either end was present near adjacent genes. Thus, we collected 1120 numbers of inter-operon IGRs for *E. coli* and 1150 numbers of inter-operon IGRs for *S. enterica* with varying sizes (Sen et al., 2022).

### Extraction of intra-operon intergenic regions

We considered all the non-zero size intra-operon IGRs for the study. The intergenic regions present between gene pairs lying under a single operon unit were focused primarily. The selection of intra-operons followed this procedure, suppose in lac operon, we found a size of 52 nucleotides of intra-operon IGRs in between *lac*Z and *lac*Y genes. Similarly, we found a size of 64 nucleotides of intra-operon IGRs in between *lac*Y and *lac*A. But in case of few operons like *tam* operon, *tam*A and *tam*B were found to be overlapped by 3 nucleotides hence we did not consider such intra-operon IGRs for the study. In supplementary tables 1 and 2, a list of intra-operon IGRs, size, GC%, location, and function is provided for *E. coli* and *S. enterica* respectively. In total, we collected 134 intra-operon IGRs for *E. coli* and 89 intra-operon IGRs in *S. enterica* from the available dataset. A size comparison boxplot was drawn between intra and inter-operon IGRs by using OriginPro 2022. OriginLab Corporation, Northampton, MA, USA. Mann-Whitney test was performed between intra-operon IGRs and inter-operon IGRs by using https://www.socscistatistics.com/tests/mannwhitney/default2.aspx website (Mann and Whitmey, 1947).

### Finding base substitutions in a sequence alignment

A hypothetical example of the detailed procedure is explained (Supplementary Table 3). Strains having ‘N’ were not considered for the study. The reference sequence was derived for each intra-operon IGRs by considering the most frequent nucleotide present at a certain position in the alignment as a reference nucleotide for that position. A similar procedure was followed for the remaining IGRs in both species. The procedure followed a methodology already established in our lab to derive the reference sequence of coding sequences (Sen et al., 2022). A python script was written to find out the spectra in both IGRs. The spectra were constructed as a total number of substitutions and normalization values. The normalization was done as per the individual substitution and dividing by the number of original nucleotides. For example, we got C→T as 20 and we found C nucleotide as 100, then the normalization was done by taking 20/100 which is 0.2.

After finding out the individual intra-operon IGRs having polymorphism in each bacterial species, the total spectra were calculated by considering only those samples having polymorphism (Supplementary Table 4). Suppose in *E. coli*, out of 5 total intra-operon IGRs we found substitution in *rpo*BC and *lac*ZY intra-operon IGRs, only thesetwo intra-operon IGRs were considered for the construction of total spectra.

### Comparison of polymorphism spectra at the intra-operon IGRs and inter-operon IGRs

After finding the intra-operon IGRs and inter-operon IGRs polymorphism spectra, we performed the comparision. We also did a comparison between Ti/Tv ratio in overall spectra and the individual nucleotide-based Ti/Tv between intra-operon IGRs and inter-operon IGRs. The overall K→M/M→K, R→Y/Y→R, and A/T→G/C or G/C→A/T bias was also observed between both the intra-operon IGRs and inter-operon IGRs. This procedure was also followed for the *S. enterica* dataset.

## Results

### Inter-operon IGRs are larger than intra-operon IGRs

In this study total 1120 number of inter-operon IGRs were considered in *E. coli*. The minimum and maximum sizes of inter-operon IGRs were 101 bp and 1384 bp respectively. Total 1150 numbers of inter-operon IGRs were considered in *S. enterica*. The minimum and maximum sizes of inter-operon IGRs were 101 bp and 1732 bp respectively. Total 134 intra-operon IGRs were studied in *E. coli* with sizes ranging from 7 to104 bp. Between *hde*A and *hde*B genes, the size of intra-operon IGRs was 104 bp in *E. coli*. Total 89 intra-operon IGRs were studied in *S. enterica* with the sizes ranging from 6 to 93 bp. Between *degQ* and *degS* genes, the size of intra-operon IGRs was 89 bp in *S. enterica*. It was distinct that intra-operon IGRs were usually smaller than inter-operon IGRs in both species (Figure 3). This is expected because intra-operon IGRs are only transcribed and not translated. So, unless used for some regulation like ribosome binding side, regulation of translation initiation, these regions will consume energy for making a transcript without any use. It is pertinent to note that intra-operon IGRs are transcribed several times depending upon the gene expression. In addition, the shorter IGRs are to avoid the rho-dependent termination. Therefore, there might be a selection for lowering the size of intra-operon IGSs. The long intra-operon IGRs such as 104 and 93 bp long indicate the possible role of translational regulation of these genes.

**Figure 3.**
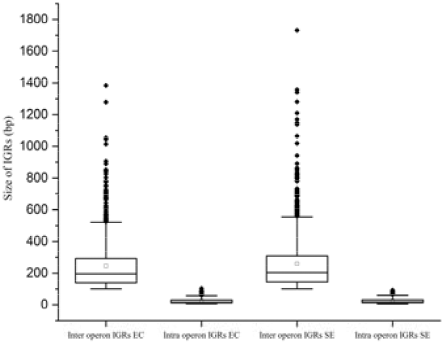
Inter-operon IGRs are larger in size than the intra-operon IGRs in *E. coli* and *S. enterica*. Legend. Box plot analysis of the size of intra-operon IGRs and inter-operon IGRs the y-axis shows the size of IGRs in numbers. The inter-operon and intra-operon IGRs box plots can be differentiated. Inter-operon IGRs show the box and whisker, even the outliers can be seen whereas intra-operon IGRs show the box and whisker right above zero. Whereas the box in inter-operon IGRs is ranged between 150-350bp. EC-*E. coli*, SE-*S. enterica. The result of the comparison between intra-operon IGRs and inter-operon IGRs was* found to be significantly different at *p <0*.*01*.

### Single nucleotide polymorphism frequency is lower at the intra-operon IGRs than the inter-operon IGRs

The polymorphism spectra in Supplementary Table 5 elucidate the total number of polymorphisms observed in inter-operon IGRs and intra-operon IGRs in *E. coli* and *S. enterica*. In *E. coli*, the inter-operon IGRs frequency was noted as 0.115, and intra-operon IGRs frequency was noted as 0.045, indicating a more than 2-fold difference between the two IGRs, whereas in *S. enterica* the inter-operon IGRs frequency was noted as 0.193 and intra-operon IGRs frequency was noted as 0.067, indicating more than a 2.5-fold difference between the two IGRs. It suggests that the intra-operon IGRs exhibit lesser nucleotide polymorphism than inter-operon IGRs contrary to the assumptions (Table 1). Along the line of the observation, out of 134 intra-operon IGRs, 77 IGRs were found to be conserved across the strains in *E. coli*. Similarly in *S. enterica*, out of 89 intra-operon IGRs, 32 IGRs were found to be conserved across the strains.

**Table 1.**
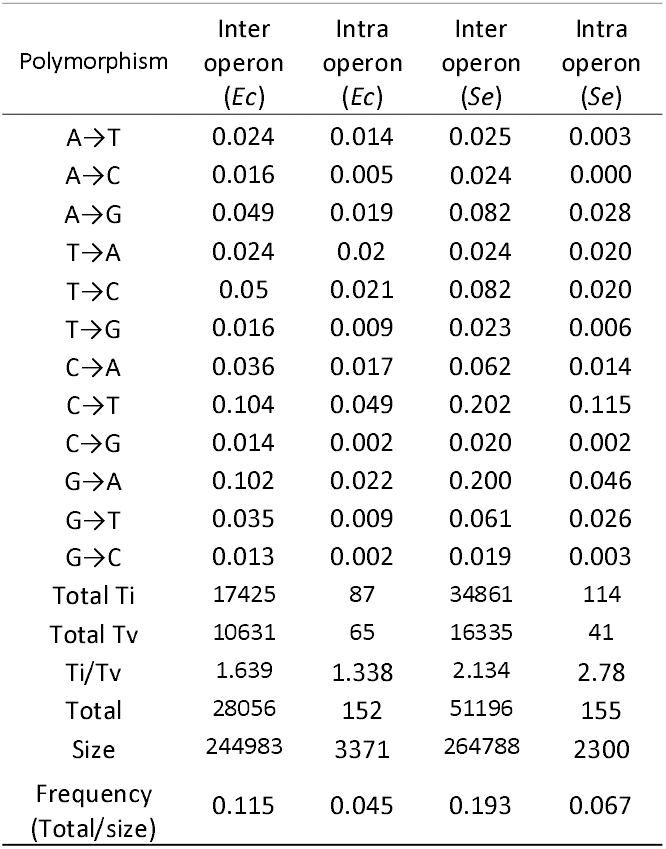
Normalized mutational spectra, Ti/Tv, and mutation frequency of *Escherichia coli (Ec) and Salmonella enterica (Se)* showing numbers and normalized values of base substitutions at inter-operon IGRs and intra-operon IGRs.

| Polymorphism | Inter operon (Ec) | Intra operon (Ec) | Inter operon (Se) | Intra operon (Se) |
| --- | --- | --- | --- | --- |
| A→T | 0.024 | 0.014 | 0.025 | 0.003 |
| A→C | 0.016 | 0.005 | 0.024 | 0.000 |
| A→G | 0.049 | 0.019 | 0.082 | 0.028 |
| T→A | 0.024 | 0.02 | 0.024 | 0.020 |
| T→C | 0.05 | 0.021 | 0.082 | 0.020 |
| T→G | 0.016 | 0.009 | 0.023 | 0.006 |
| C→A | 0.036 | 0.017 | 0.062 | 0.014 |
| C→T | 0.104 | 0.049 | 0.202 | 0.115 |
| C→G | 0.014 | 0.002 | 0.020 | 0.002 |
| G→A | 0.102 | 0.022 | 0.200 | 0.046 |
| G→T | 0.035 | 0.009 | 0.061 | 0.026 |
| G→C | 0.013 | 0.002 | 0.019 | 0.003 |
| Total Ti | 17425 | 87 | 34861 | 114 |
| Total Tv | 10631 | 65 | 16335 | 41 |
| Ti/Tv | 1.639 | 1.338 | 2.134 | 2.78 |
| Total | 28056 | 152 | 51196 | 155 |
| Size | 244983 | 3371 | 264788 | 2300 |
| Frequency (Total/size) | 0.115 | 0.045 | 0.193 | 0.067 |

### Comparative polymorphism spectra analysis of inter-operon IGRs and intra-operon IGRs

The nucleotide polymorphism spectra in the inter-operon IGRs of *E. coli* and *S. enterica* were known from the earlier research in the laboratory (Sen et al. 2022). In brief, transition polymorphisms such as C→T and G→A were two times more frequent than T→C and A →G. In case of Tv polymorphisms, G→T and C→A were more frequent than the others. The complementary polymorphisms were of similar values (Sen et al. 2022). In this study we found out intra-operon IGRs complementary polymorphisms were of different values, unlike the inter-operon IGRs. The transition polymorphism C→T value was 0.049 while the other transition polymorphisms such as G→A, A→G, T→C were 0.022. 0.019 and 0.021, respectively. The highest Tv was obtained in T→A with a value of 0.020, which was almost like three other Ti spectra A→G, T→C, and G→A. Among other Tv frequency values, C→A was observed to be the second highest Tv with a value of 0.017 followed by A→T of 0. 014. The complementary substitutions between A→T (0.014) and T→A (0.020), a 1.4-fold value difference was observed, likewise between A→C (0.005) and T→G (0.009) also a fold difference of 1.8 was observed. The fold difference between C→A (0.017) and G→T (0.009) a 2-fold difference was observed. The inter-operon IGRs and intra-operon IGRs polymorphism spectra were found to be significantly different at p<.05 (Figure 4).

**Figure 4.**
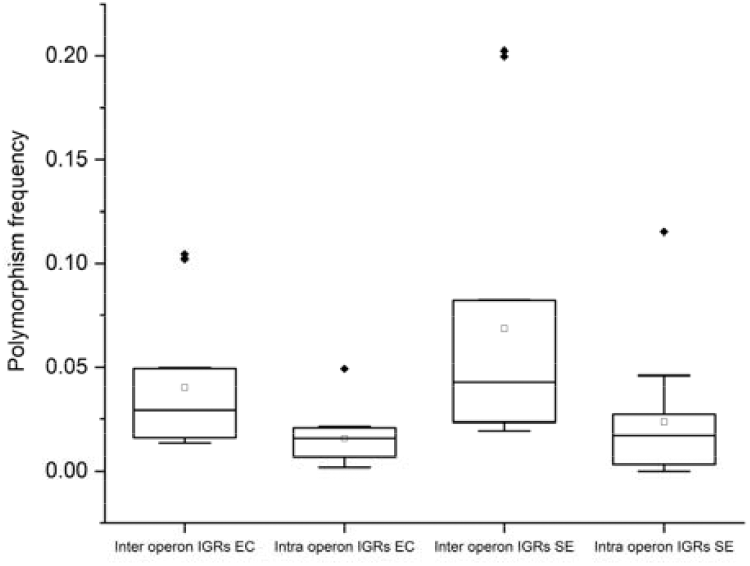
The inter-operon IGRs polymorphism values were significantly more than that of intra-operon IGRs in both the bacteria. Legend. Boxplot showing polymorphism frequency/normalization values in y-axis. EC-*E. coli* and SE-*S. enterica*. the inter-operon IGRs polymorphism values are significantly more than intra-operon IGRs in both the species. In Mann Whitney u test also the values between both IGRs in both species were found to be significantly different at p<0.05

Similarly in *S. enterica* in brief, transition polymorphisms such as C→T and G→A were more than two times frequent than T→C and A →G. In case of Tv polymorphisms, G→T and C→A were more frequent than the others. The complementary polymorphisms were of similar values (Sen et al. 2022). In this study we found out intra-operon IGRs complementary polymorphisms were of different values, unlike the inter-operon IGRs. The transition polymorphism C→T value was 0.115 while the other transition polymorphisms such as G→A, A→G, T→C were 0.046. 0.028 and 0.020, respectively. The highest Tv was obtained in T→A with a value of 0.020, which was almost like three other Ti spectra A→G, T→C, and G→A. Among other Tv frequency values, G→T was observed to be the second highest Tv with a value of 0.026 followed by T→A of 0. 020. The complementary substitutions between A→T (0.003) and T→A (0.020), a six-fold value difference was observed, likewise between G→T (0.026) and C→A (0.014) also a fold difference of 1.8 was observed.

### Ti/Tv values at inter-operon and intra-operon IGRs in these bacteria

In *E. coli*, the ratio of Ti/Tv was observed as 1.639 at inter-operon IGRs and 1.338 at intra-operon IGRs (Table 1). We then found out Ti/Tv at individual nucleotides for inter-operon IGRs. For Adenine, A→G Ti was compared with combined A→T and A→C Tv. A→G value was 0.049 and the combined A→T and A→C values was 0.040, hence the ratio of Ti/Tv for A was 1.22 (Table 2). The Ti/Tv ratio T, C, and G were obtained as 1.25, 2.93, and 2.13 respectively. The Ti/Tv in case of G and C were distinctly higher than that for A and T. So, transition was more frequent than transversion in a genome even at individual nucleotide level. In intra-operon IGRs the individual nucleotide-based Ti/Tv ratio at A, T, C, and G were obtained as 1.00, 0.74, 2.58 and 1.9 respectively (Table 2). The ratio at T (0.074) indicated the existence of higher Tv values in T, as explained above T→A Tv value exhibited unexpectedly equal values with T→C. Hence the polymorphism pattern relating to T nucleotide was observed to be different.

**Table 2.** individual nucleotide-based Ti/Tv at inter and inter-operon IGRs in both the species along with GC%.

| Species | IGRs | Individual nucleotide-based Ti/Tv |  |  |  | GC% |
| --- | --- | --- | --- | --- | --- | --- |
|  |  | A | T | C | G |  |
| *E. coli* | Inter operon | 1.22 | 1.25 | 2.93 | 2.13 | 40.3 |
|  | Intra operon | 1 | 0.74 | 2.58 | 1.9 | 44.8 |
| *S. enterica* | Inter operon | 1.69 | 1.74 | 2.47 | 2.5 | 41.6 |
|  | Intra operon | 10 | 0.77 | 7 | 1.56 | 47.6 |

In *S. enterica*, the ratio of Ti/Tv was observed as 2.134 at inter-operon IGRs and 2.780 at intra-operon IGRs (Table 1). Ti/Tv at individual nucleotides were calculated like *E. coli*. A→G value was 0.082 and the combined A→T and A→C values was 0.049, hence the ratio of Ti/Tv for A was 1.69 (Table 2). The Ti/Tv ratio T, C, and G were obtained as 1.74, 2.47, and 2.50 respectively. The Ti/Tv in case of G and C were distinctly higher than that for A and T similar to *E. coli*. In intra-operon IGRs the individual nucleotide-based Ti/Tv ratio at A, T, C, and G were obtained as 10.00, 0.77, 7.00 and 1.56 respectively (Table 2). The ratio at T (0.077) indicated the existence of higher Tv values in T, as explained above T→A Tv value exhibited unexpectedly equal values with T→C. This similar observation was also seen in case of *E. coli*.

The comparative individual nucleotide-based substitutions between inter and intra-operon IGRs in *S. enterica* resembled the results in *E. coli* (Table 2). Ti/Tv ratio at T in intra-operon IGRs was found to be 0.77 which indicated the unusual T→A spectra value has similar values of T→C.

### Intra-operon polymorphism is biased towards purine and keto nucleotides

The comparative study of different biases (RY, KM, AT/GC) revealed that there was difference between inter-operon IGRs and intra-operon IGRs in *E. coli* (Table 3). In inter-operon IGRs, the R→Y(Pu→Py) substitution was observed as 0.088, whereas Y→R for the same was observed as 0.089. But in case of intra-operon IGRs the R→Y was found to be 0.030 and Y→R was found to be 0.048, a 1.6-fold difference was observed in *E. coli*, which was showing a higher proportion of Pu biased polymorphism in intra-operon IGRs. Similarly,while analyzing K→M and M→K (Keto/Amino) biased polymorphism we found similar values in case of inter-operon IGRs for both. But in case of intra-operon IGRs, K→M was observed to be 0.065 and M→K was observed to be 0.084 indicating 1.2-fold higher Keto biased polymorphism in intra-operon IGRs. When we calculated the difference between AT and GC biased polymorphism in both IGRs, intra-operon IGRs were found as 0.044 and inter-operon IGRs as 0.146, which recommended a higher AT biased mutation in inter-operon IGRs. Similar observations were also found in *S. enterica* regarding the nucleotide polymorphism spectra at inter-operon IGRs (Table 1). Analogous to *E. coli*, the keto and Purine biases were also observed in intra-operon IGRs, as well as higher AT biases were also observed in inter-operon IGRs (Table 3).

**Table 3.** RY, KM, AT/GC biases present between inter-operon IGRs and intra-operon IGRs in *E. coli* and *S. enterica*.

| Species and IGRs | R→Y | Y→R | K→M | M→K | A/T→G/C | G/C→A/T | Difference<br>(A/T)-(G/C) |
| --- | --- | --- | --- | --- | --- | --- | --- |
| *E. coli* Inter operon | 0.088 | 0.089 | 0.189 | 0.191 | 0.131 | 0.277 | 0.146 |
| *E. coli* Intra operon | 0.030 | 0.048 | 0.065 | 0.084 | 0.054 | 0.097 | 0.044 |
| *S. enterica* Inter operon | 0.128 | 0.129 | 0.325 | 0.330 | 0.211 | 0.525 | 0.313 |
| *S. enterica* Intra operon | 0.032 | 0.042 | 0.089 | 0.149 | 0.054 | 0.202 | 0.148 |

Hence the frequency comparison, individual nucleotide-based Ti/Tv ratio, skew biased comparison and overall spectra suggested that intra-operon IGRs have a different polymorphism than inter-operon IGRs which possibly explains the role of mutation and/or selection during transcriptional mutagenesis on intra-operon IGRs.

## Discussion

Since intra-operon IGRs undergo both replication and transcription, the probability of getting elevated numbers of polymorphism was expected due to the cumulative effect of both replication and transcription in comparison to the inter-operon IGRs. But our observation suggests that it might not always be true considering the influence of selection and transcription-coupled DNA repair. Despite replication and transcription, the intra-operon IGRs were observed to be more conserved across the genomes of different strains in *E. coli* and *S. enterica*. It will be very interesting to compare the intra-operon IGRs spectra in high and low-expression genes in the future as highly expressed genes undergo single-stranded separation more frequently than low-expressed genes.

Intra-operon IGRs are under different selection mechanisms than the inter-operon IGRs as follows: transcription coupled repair; rho-dependent termination; and ribosome binding site for translation initiation. These might attribute for the low frequency of polymorphism in these regions. The increase in purine at intra-operon IGRs in favor of ribosome binding site for the downstream gene and decreases the probability of a rho-dependent termination site for the upstream gene (Bogden et al. 1999). The increase in keto nucleotides might be favoring any need of secondary structure (Sen et al. 2022).

Usually, transitions are more frequently selected over transversion in a genome. But when a region of a chromosome has an almost equal proportion of transition and transversion relating to one nucleotide, the role of mutation and/or selection cannot be denied. Such transversions are allowable only when it has a benefic effect on the fitness of an organism. For translational efficacy, the prokaryotic ribosome binding sites (RBS) near the Shine-Dalgarno (SD) sequences are selected for Purine-rich nucleotides (Shine and Dalgarno 1974; Omotajo et al. 2015). Hence the frequent T→A unusual base substitutions at intra-operon IGRs might have arisen or been selected to facilitate in translation of the downstream gene. Future research will shed more light on these aspects of the base substitutions in intra-operon IGRs and how differently they are functioning from inter-operon IGRs even in other species.

The comparative study of inter-operon IGR and intra-operon IGR might be investigated in more detailed to develop polymorphisms signatures to discriminate between these two regions in a genome, which might be helpful for genome annotation and defining operon in bacterial genome. Therefore, the observation of lower polymorphism frequency, high T→A transversion, polymorphism bias for purine and keto nucleotides might be useful in predicting inter-operon *vs* intra-operon IGRs.

## Acknowledgments

PKB is grateful to Tezpur University for the institutional fellowship. PS is grateful to UGC, GoI New Delhi, for the JRF. RA is thankful for the JRF fellowship from the DBT grant (BT/511/NE/TBP/2013) and (BT/403/NE/U-Excel/2013). SSS and SKR are grateful to DBT, GoI, for the Computational Biology and Bioinformatics Center at Tezpur University.

## Authors’ Contributions

Conceptualization: PKB, SSS, SKR; Methodology: PKB, PS, SSS and SKR; Writing, review, and editing: PKB, PS, RA, SSS and SKR; Supervision: SSS and SKR. All authors endorsed the manuscript.

