## Supplementary_materials for "The transcribed intergenic regions exhibit lower frequency of nucleotide polymorphism than the untranscribed intergenic regions in the genomes of *Escherichia coli and Salmonella enterica*"

Supplementary_material

**Supplementary table 1. *E. coli* intra operon IGRs list with coordinate details of adjacent genes, size and GC % of IGRs and biological functional information on individual IGRs^[1]^**

| Sl no. | Intra operon | Location | | Size | GC% | Information |
| --- | --- | --- | --- | --- | --- | --- |
|  |  | Gene-1 | Gene-2 |  |  |  |
| 1 | *hdeAB* | 3655966..3656304 | 3656408..3656740 | 104 | 37.50 | Acid stress chaperone unit |
| 2 | *tnaAB* | 3888730..3890145 | 3890236..3891483 | 91 | 39.33 | Unit of trp operon |
| 3 | *degQS* | 3380743..3382110 | 3382200..3383267 | 90 | 51.14 | serine endoprotease |
| 4 | *dnaKJ* | 12163..14079 | 14090..14167 | 87 | 47.13 | Chaperone |
| 5 | *cadBA* | 4356470..4358617 | 4358697..4360031 | 80 | 42.50 | Lysine metabolism |
| 6 | *rpoBC* | 4181245..4185273 | 4185350..4189573 | 77 | 54.55 | RNA polymerase subunits |
| 7 | *guaBA* | 2630958..2632535 | 2632604..2634070 | 69 | 44.93 | nucleotide metabolism |
| 8 | *lacYA* | 361249..361860 | 361926..363179 | 64 | 34.92 | lactose metabolism |
| 9 | *manXY* | 1902048..1903019 | 1903082..1903882 | 63 | 46.67 | mannose metabolism |
| 10 | *nagBA* | 701603..702751 | 702811..703611 | 60 | 50.00 | Glucosamine metabolism |
| 11 | *nhaAR* | 17489..18655 | 18721..19620 | 60 | 44.64 | Sodium antiporter |
| 12 | *xapAB* | 2522729..2523985 | 2524045..2524878 | 58 | 44.64 | nucleotide metabolism |
| 13 | *creCD* | 4636696..4638120 | 4638178..4639530 | 56 | 48.08 | Histidine kinase regulation |
| 14 | *deoBD* | 4619603..4620826 | 4620883..4621602 | 56 | 63.27 | nucleotide metabolism |
| 15 | *lacZY* | 361926..363179 | 363231..366305 | 52 | 38.00 | lactose metabolism |
| 16 | *atpAG* | 3917402..3918265 | 3918316..3919857 | 51 | 48.98 | ATP synthase |
| 17 | *deoAB* | 4618229..4619551 | 4619603..4620826 | 50 | 54.35 | nucleotide metabolism |
| 18 | *fhuAC* | 167484..169727 | 169778..170575 | 49 | 36.96 | iron metabolism |
| 19 | *livKH* | 3595477..3596403 | 3596451..3597560 | 46 | 41.30 | Amino acid ABC transporter |
| 20 | *hicAB* | 1509286..1509462 | 1509487..1509924 | 46 | 57.78 | toxin-antitoxin |
| 21 | *fliAZ* | 2000473..2001024 | 2001070..2001789 | 46 | 38.64 | flagella biosynthesis |
| 22 | *nrfAB* | 4287764..4289200 | 4289245..4289811 | 45 | 59.09 | cytochrome metabolism |
| 23 | *groSL* | 4370688..4370981 | 4371025..4372671 | 44 | 39.53 | cochaperonin |
| 24 | *eutKR* | 2554130..2555183 | 2555228..2555729 | 44 | 46.55 | ethanolamine utilization protein |
| 25 | *ptsHI* | 2533764..2534021 | 2534066..2535793 | 43 | 43.14 | phosphocarrier protein |
| 26 | *nuoFG* | 2397439..2400165 | 2400218..2401555 | 41 | 35.90 | NADH dehydrogenase subunit |
| 27 | *csgBA* | 1103951..1104406 | 1104447..1104902 | 39 | 50.00 | curlin subunit |
| 28 | *prpDE* | 351215..352666 | 352706..354592 | 38 | 27.03 | methylcitrate metabolsim |
| 29 | *agaBC* | *3284170..3284646* | *3284685..3285488* | 37 | 39.39 | N-acetylgalactosamine transporter |
| 30 | *rpsN/rpsH* | 3446153..3446545 | 3446579..3446884 | 34 | 41.18 | ribosomal units |
| 31 | *rpsJ/rplC* | 3452297..3452926 | 3452959..3453270 | 33 | 42.42 | ribosomal units |
| 32 | *prpCD* | *350012..351181* | *351215..352666* | 32 | 40.63 | methylcitrate metabolsim |
| 33 | *tdcDE* | 3260124..3262418 | 3262452..3263660 | 32 | 25.00 | propionic acid metabolism |
| 34 | *fucPI* | 2934235..2935551 | 2935584..2937359 | 31 | 32.26 | L-fucose metabolism |
| 35 | *caiTA* | 39244..40386 | 40417..41931 | 31 | 34.38 | gamma-butyrobetaine antiporter |
| 36 | *marAB* | 1619574..1619957 | 1619989..1620207 | 30 | 43.33 | transcriptional regulators |
| 37 | *atoEB* | 2324756..2326078 | 2326109..2327293 | 29 | 44.83 | fatty acid metabolism |
| 38 | *aceBA* | 4215478..4217079 | 4217109..4218413 | 28 | 46.43 | TCA cycle |
| 39 | *fucAO* | 2931865..2933013 | 2933041..2933688 | 28 | 39.29 | L-fucose fermentation |
| 40 | *lsrBF* | 1605051..1606073 | 1606100..1606975 | 27 | 37.04 | Autoinducer-2 ABC transporter |
| 41 | *cmtBA* | 3077471..3078859 | 3078887..3079330 | 26 | 34.62 | mannitol metabolism |
| 42 | *atpGD* | 3915993..3917375 | 3917402..3918265 | 25 | 34.62 | ATP synthase |
| 43 | *ycbKL* | *1112718..1113267* | *1113291-1113940* | 24 | 62.50 | periplasmic protein |
| 44 | *tdcCD* | 3262452..3263660 | 3263686..3265017 | 24 | 45.83 | Thr metabolism |
| 45 | *gcvTH* | 3049160..3049549 | 3049573..3050667 | 24 | 25.00 | Gly metabolism |
| 46 | *flgDE* | 1131854..1132549 | 1132574..1133782 | 23 | 47.83 | flagella biosynthesis |
| 47 | *minCD* | 1224549..1225361 | 1225385..1226080 | 23 | 30.43 | cell division protein |
| 48 | *yphED* | 1458264..1459052 | 1459099..1460481 | 23 | 43.48 | ABC transporter permease |
| 49 | *kdpAB* | 724988..727036 | 727059..728732 | 23 | 34.78 | potassium-transporting ATPase |
| 50 | *moaAB* | 817044..818033 | 818055..818567 | 22 | 40.91 | molybdenum biosynthesis |
| 51 | *hemDX* | 3987885..3989066 | 3989088..3989828 | 22 | 40.91 | uroporphyrinogen biosynthesis |
| 52 | *tdcBC* | 3263686..3265017 | 3265039..3266028 | 22 | 31.82 | Thr metabolism |
| 53 | *yphFE* | 2676850..2678361 | 2678384..2679367 | 21 | 61.90 | ABC transporter |
| 54 | *rpmBG* | 3811250..3811417 | 3811438..3811674 | 21 | 28.57 | 50S ribosomal protein |
| 55 | *cyoAB* | 448650..450641 | 450663..451610 | 20 | 45.00 | cytochrome metabolism |
| 56 | *cheBY* | 1967048..1967437 | 1967452..1968501 | 20 | 40.00 | chemotaxis |
| 57 | *marRA* | 1619120..1619554 | 1619574..1619957 | 20 | 35.00 | transcriptional regulators |
| 58 | *clpSA* | 922913..923233 | 923264..925540 | 19 | 52.63 | ATP-dependent protease |
| 59 | *eutBC* | 2556410..2557297 | 2557318..2558679 | 19 | 47.37 | ethanolamine biosynthesis |
| 60 | *atpDC* | 3915553..3915972 | 3915993..3917375 | 19 | 52.63 | ATP synthase |
| 61 | *flgEF* | 1132574..1133782 | 1133802..1134557 | 18 | 61.11 | flagella biosynthesis |
| 62 | *carAB* | 29651..30799 | 30817..34038 | 18 | 44.44 | carbamoyl-phosphate synthetase |
| 63 | *hycDE* | 2844762..2846471 | 2846489..2847412 | 18 | 33.33 | formate hydrogenlyase |
| 64 | *hyaBC* | 1033254..1035047 | 1035066..1035773 | 17 | 47.06 | hydrogenase metabolism |
| 65 | *fliIJ* | 2016554..2017927 | 2017946..2018389 | 17 | 58.82 | flagella biosynthesis |
| 66 | *emrAB* | 2811427..2812599 | 2812616..2814154 | 17 | 58.82 | MDR protein |
| 67 | *feoAB* | 3540163..3540390 | 3540407..3542728 | 17 | 35.29 | iron metabolism |
| 68 | *cusFB* | 597131..597463 | 597479..598702 | 16 | 43.75 | Cu/Ag export |
| 69 | *artPI* | 902257..902988 | 903006..903734 | 16 | 37.50 | Arg metabolism |
| 70 | *mglAC* | 2236743..2237753 | 2237769..2239289 | 16 | 56.25 | carbohydrate metabolism |
| 71 | *rplW/rplB* | 3450543..3451364 | 3451382..3451684 | 16 | 43.75 | ribosomal units |
| 72 | *rplV/rpsC* | 3449182..3449883 | 3449901..3450233 | 16 | 50.00 | ribosomal units |
| 73 | *cydAB* | 771458..773026 | 773042..774181 | 16 | 50.00 | cytochrome |
| 74 | *speED* | 134788..135582 | 135598..136464 | 16 | 31.25 | spermidine synthase |
| 75 | *rcsDB* | 2313488..2316160 | 2316177..2316827 | 15 | 40.00 | phosphorelay system |
| 76 | *rplB/rpsS* | 3450248..3450526 | 3450543..3451364 | 15 | 40.00 | ribosomal units |
| 77 | *fabHD* | 1148759..1149712 | 1149728..1150657 | 14 | 21.43 | oxoacyl biosynthesis |
| 78 | *nuoAB* | 2403951..2404613 | 2404629..2405072 | 14 | 57.14 | NADH dehydrogenase |
| 79 | *lepAB* | 2704335..2705309 | 2705325..2707124 | 14 | 57.14 | 30s ribosomoal biosynthesis |
| 80 | *rplM/rpsI* | 3377815..3378207 | 3378223..3378651 | 14 | 50.00 | ribosomal units |
| 81 | *potAB* | 1183617..1184474 | 1184458..1185594 | 14 | 35.71 | ABC transporter |
| 82 | *fixAB* | 42403..43173 | 43188..44129 | 13 | 53.85 | electron transfer flavoprotein |
| 83 | *citCD* | 650487..650783 | 650798..651856 | 13 | 30.77 | citrate lyase synthetase |
| 84 | *sucAB* | 758706..761507 | 761522..762739 | 13 | 30.77 | 2-oxoglutarate dehydrogenase |
| 85 | *oppBC* | 1302899..1303819 | 1303834..1304742 | 13 | 61.54 | oligopeptide transporter |
| 86 | *araGH* | 1982554..1983540 | 1983555..1985069 | 13 | 38.46 | arabisone operon |
| 87 | *nuoHI* | 2395908..2396450 | 2396465..2397442 | 13 | 53.85 | NADH dehydrogenase |
| 88 | *rplR/rpsE* | 3444726..3445229 | 3445244..3445597 | 13 | 46.15 | ribosomal units |
| 89 | *rpsS/rplV* | 3449901..3450233 | 3450248..3450526 | 13 | 53.85 | ribosomal units |
| 90 | *rplx/rplE* | 3446899..3447438 | 3447453..3447767 | 13 | 46.15 | ribosomal units |
| 91 | *rplE/rpsN* | 3446579..3446884 | 3446899..3447438 | 13 | 38.46 | ribosomal units |
| 92 | *atpFH* | 3919870..3920403 | 3920418..3920888 | 13 | 76.92 | ATP synthase |
| 93 | *fdoGH* | 4081857..4082759 | 4082772..4085186 | 13 | 46.15 | formate dehydrogenase |
| 94 | *malFG* | 4242626..4243516 | 4243531..4245075 | 13 | 30.77 | maltose metabolism |
| 95 | *pyrBI* | 4470986..4471447 | 4471460..4472395 | 13 | 46.15 | aspartate biosynthesis |
| 96 | *tauAB* | 385232..386194 | 386207..386974 | 13 | 61.54 | Taurine metabolism |
| 97 | *ubiCA* | 4252506..4253003 | 4253016..4253888 | 13 | 23.08 | chorismate pyruvate lyase |
| 98 | *betBA* | 325577..327247 | 327261..328733 | 12 | 41.67 | choline, oxygen, and osmotic stress |
| 99 | *entEB* | 626070..627680 | 627694..628551 | 12 | 50.00 | 2,3-dihydroxybenzoate-AMP ligase |
| 100 | *gabDT* | 2791273..2792721 | 2792735..2794015 | 12 | 58.33 | succinate-semialdehyde dehydrogenase |
| 101 | *cusBA* | 597479..598702 | 598714..601857 | 12 | 41.67 | Cu/Ag export |
| 102 | *paaAB* | 1453927..1454856 | 1454868..1455155 | 12 | 33.33 | monooxygenase subunit |
| 103 | *fabDG* | 1149728..1150657 | 1150670..1151404 | 11 | 27.27 | S-malonyltransferase |
| 104 | *manYZ* | 1903082..1903882 | 1903895..1904746 | 11 | 63.64 | mannose metabolism |
| 105 | *rpsC/rplP* | 3448759..3449169 | 3449182..3449883 | 11 | 72.73 | ribosomal units |
| 106 | *rpsH/rplF* | 3445607..3446140 | 3446153..3446545 | 11 | 45.45 | ribosomal units |
| 107 | *atpHA* | 3918316..3919857 | 3919870..3920403 | 11 | 72.73 | ATP synthase |
| 108 | *bcsAB* | 3690268..3692571 | 3692618..3695236 | 11 | 45.45 | cellulose synthase |
| 109 | *araBA* | 66835..68337 | 68348..70048 | 11 | 45.45 | arabisone operon |
| 110 | *frdCD* | 4379007..4379366 | 4379377..4379772 | 11 | 63.64 | fumarate reductase |
| 111 | *citEF* | 648039..649571 | 649582..650490 | 11 | 36.36 | citrate metabolism |
| 112 | *proBA* | 260388..261491 | 261503..262756 | 10 | 70.00 | gamma-glutamyl kinase |
| 113 | *cyoDE* | 446815..447705 | 447717..448046 | 10 | 70.00 | cytochrome metabolism |
| 114 | *trpCB* | 1317222..1318415 | 1318427..1319785 | 10 | 40.00 | trp operon |
| 115 | *flgCD* | 1131438..1131842 | 1131854..1132549 | 10 | 50.00 | flagella biosynthesis |
| 116 | *oppCD* | 1303834..1304742 | 1304754..1305767 | 10 | 20.00 | oligopeptide transport system |
| 117 | *nuoIJ* | 2395342..2395896 | 2395908..2396450 | 10 | 70.00 | NADH dehydrogenase |
| 118 | *acrEF* | 3413864..3415021 | 3415033..3418137 | 10 | 40.00 | acriflavin resistance protein |
| 119 | *glnLG* | 4053869..4055278 | 4055290..4056339 | 10 | 50.00 | nitrogen regulation |
| 120 | *malPQ* | 3547986..3550070 | 3550080..3552473 | 10 | 60.00 | maltodextrin phosphorylase |
| 121 | *glyQS* | 3722328..3724397 | 3724407..3725318 | 10 | 70.00 | glycyl-tRNA synthetase |
| 122 | *dadAX* | 1237571..1238869 | 1238879..1239949 | 10 | 50.00 | amino acid metabolism |
| 123 | *fadBA* | 4027609..4028772 | 4028782..4030971 | 10 | 50.00 | fatty acid metabolism |
| 124 | *codBA* | 354922..356181 | 356171..357454 | 10 | 40.00 | cytosine metabolism |
| 125 | *leuCD* | 78848..79453 | 79464..80864 | 9 | 54.55 | isopropylmalate dehydratase subunit |
| 126 | *flgHI* | 1135564..1136262 | 1136274..1137371 | 9 | 66.67 | flagella biosynthesis |
| 127 | *cheYZ* | 1966393..1967037 | 1967048..1967437 | 9 | 55.56 | chemotaxis |
| 128 | *srlEB* | 2826392..2827351 | 2827362..2827733 | 9 | 44.44 | glucitol/sorbitol-specific |
| 129 | *accBC* | 3405436..3405906 | 3405917..3407266 | 9 | 66.67 | acetyl-CoA carboxylase carboxyl carrier |
| 130 | *rplC/rplD* | 3451681..3452286 | 3452297..3452926 | 9 | 44.44 | ribosomal units |
| 131 | *rplN/rplX* | 3447453..3447767 | 3447778..3448149 | 9 | 55.56 | ribosomal units |
| 132 | *entCE* | 624885..626060 | 626070..627680 | 8 | 75.00 | isochorismate synthase |
| 133 | *rplF/rplR* | 3445244..3445597 | 3445607..3446140 | 8 | 37.50 | ribosomal units |
| 134 | *nagAC* | 700374..701594 | 701603..702751 | 7 | 28.57 | N-acetylglucosamine biosynthesis |

**Supplementary table 2. *S. enterica* intra operon IGRs list with coordinate details of adjacent genes, size and GC % of IGRs and biological functional information on individual IGRs^[2]^**

| Sl no. | Intra operon | Location | | Size | GC% | Information |
| --- | --- | --- | --- | --- | --- | --- |
|  |  | Gene-1 | Gene-2 |  |  |  |
| 1 | *degQS* | 3538553..3539920 | 3540013..3541083 | 93 | 51.61 | serine endoprotease |
| 2 | *dnaKJ* | 13520..13594 | 11593..13509 | 86 | 51.16 | Chaperone |
| 3 | *cadBA* | 2722916..2724247 | 2724330..2726474 | 83 | 53.01 | Lysine metabolism |
| 4 | *rpoBC* | 4388659..4392687 | 4392764..4396987 | 77 | 53.25 | RNA polymerase subunits |
| 5 | *guaBA* | 2647184..2648761 | 2648831..2650297 | 70 | 45.71 | nucleotide metabolism |
| 6 | *nhaAR* | 46190..47356 | 47424..48317 | 68 | 51.47 | Sodium antiporter |
| 7 | *tdcCD* | 3431184..3432392 | 3432458..3433789 | 66 | 66.67 | Thr metabolism |
| 8 | *mreBC* | 3565538..3566590 | 3566655..3567698 | 65 | 53.85 | dynamic cytoskeletal protein |
| 9 | *livKH* | 3754280..3755206 | 3755266..3756375 | 60 | 53.33 | Amino acid ABC transporter |
| 10 | *fliAZ* | 2027647..2028198 | 2028257..2028976 | 59 | 54.24 | flagella biosynthesis |
| 11 | *creCD* | 4866890..4868314 | 4868372..4869721 | 58 | 48.28 | Histidine kinase regulation |
| 12 | *xapAB* | 2557213..2558469 | 2558524..2559357 | 55 | 45.45 | nucleotide metabolism |
| 13 | *deoAB* | 4844302..4845624 | 4845676..4846899 | 52 | 32.69 | nucleotide metabolism |
| 14 | *atpAG* | 439064..441352 | 441364..441828 | 51 | 60.78 | ATP synthase |
| 15 | *fhuAC* | 223732..225921 | 225970..226767 | 49 | 53.06 | iron metabolism |
| 16 | *groSL* | 4596762..4597055 | 4597099..4598745 | 44 | 40.91 | cochaperonin |
| 17 | *csgBA* | 1225125..1225520 | 1225525..1226175 | 42 | 40.48 | curlin subunit |
| 18 | *prpCD* | 459491..460660 | 460700..462151 | 40 | 35.00 | methylcitrate metabolsim |
| 19 | *caiTA* | 85510..86652 | 86687..88204 | 35 | 34.29 | gamma-butyrobetaine antiporter |
| 20 | *rpsN/rpsH* | 3612512..3612904 | 3612938..3613243 | 34 | 41.18 | ribosomal units |
| 21 | *tdcDE* | 3428856..3431150 | 3431184..3432392 | 34 | 35.29 | propionic acid metabolism |
| 22 | *rpsJ/rplC* | 3618655..3619284 | 3619317..3619628 | 33 | 45.45 | ribosomal units |
| 23 | *aceBA* | 4424748..4426349 | 4426381..4427685 | 32 | 40.63 | TCA cycle |
| 24 | *citCD* | 68712..69680 | 69710..70003 | 30 | 36.67 | citrate lyase synthetase |
| 25 | *lsrBF* | 4309855..4310877 | 4310906..4311781 | 29 | 34.48 | Autoinducer-2 ABC transporter |
| 26 | *atpGD* | 4096987..4098369 | 4098396..4099259 | 27 | 37.04 | ATP synthase |
| 27 | *flgDE* | 1254883..1255581 | 1255608..1256819 | 27 | 48.15 | flagella biosynthesis |
| 28 | *gcvTH* | 3206432..3206821 | 3206847..3207941 | 26 | 26.92 | Gly metabolism |
| 29 | *fliIJ* | 2043407..2044777 | 2044799..2045242 | 22 | 54.55 | flagella biosynthesis |
| 30 | *moaAB* | 909257..910279 | 910301..910813 | 22 | 31.82 | molybdenum biosynthesis |
| 31 | *flgEF* | 1255608..1256819 | 1256840..1257595 | 21 | 57.14 | flagella biosynthesis |
| 32 | *kdpAB* | 809098..811146 | 811167..812846 | 21 | 42.86 | potassium-transporting ATPase |
| 33 | *rpmBG* | 3944915..3945082 | 3945103..3945339 | 21 | 28.57 | 50S ribosomal protein |
| 34 | *speED* | 194195..194989 | 195010..195870 | 21 | 33.33 | spermidine synthase |
| 35 | *carAB* | 75881..77029 | 77048..80275 | 19 | 42.11 | carbamoyl-phosphate synthetase |
| 36 | *feoAB* | 3686962..3687189 | 3687208..3689526 | 19 | 26.32 | iron metabolism |
| 37 | *artPI* | 998942..999673 | 999691..1000419 | 18 | 44.44 | Arg metabolism |
| 38 | *dsdXA* | 4025973..4027310 | 4027328..4028650 | 18 | 38.89 | D-serine metabolism |
| 39 | *hycDE* | 2986879..2988588 | 2988606..2989529 | 18 | 33.33 | formate hydrogenlyase |
| 40 | *rplV/rpsC* | 3615540..3616241 | 3616259..3616591 | 18 | 50.00 | ribosomal units |
| 41 | *rplW/rplB* | 3616901..3617722 | 3617740..3618042 | 18 | 44.44 | ribosomal units |
| 42 | *emrAB* | 2954763..2955935 | 2955952..2957490 | 17 | 58.82 | MDR protein |
| 43 | *fucAO* | 3051020..3051796 | 3051801..3052439 | 17 | 58.82 | L-fucose fermentation |
| 44 | *lepAB* | 2750161..2751135 | 2751152..2752951 | 17 | 52.94 | 30s ribosomoal biosynthesis |
| 45 | *rcsDB* | 2390974..2393643 | 2393660..2394310 | 17 | 52.94 | phosphorelay system |
| 46 | *rplB/rpsS* | 3616606..3616884 | 3616901..3617722 | 17 | 35.29 | ribosomal units |
| 47 | *cydAB* | 848339..849907 | 849923..851062 | 16 | 50.00 | cytochrome |
| 48 | *fabHD* | 1273112..1274065 | 1274081..1275010 | 16 | 25.00 | oxoacyl biosynthesis |
| 49 | *mglAC* | 2307594..2308604 | 2308620..2310140 | 16 | 56.25 | carbohydrate metabolism |
| 50 | *potAB* | 1306983..1307846 | 1307830..1308966 | 16 | 31.25 | ABC transporter |
| 51 | *aceEF* | 176242..178905 | 178920..180809 | 15 | 26.67 | pyruvate dehydrogenase |
| 52 | *atpFH* | 4100864..4101397 | 4101412..4101882 | 15 | 73.33 | ATP synthase |
| 53 | *malFG* | 4469458..4470348 | 4470363..4471907 | 15 | 33.33 | maltose metabolism |
| 54 | *nuoHI* | 2454029..2454571 | 2454586..2455563 | 15 | 53.33 | NADH dehydrogenase |
| 55 | *oppBC* | 1830158..1831066 | 1831081..1832001 | 15 | 60.00 | oligopeptide transporter |
| 56 | *rplR/rpsE* | 3611085..3611588 | 3611603..3611956 | 15 | 46.67 | ribosomal units |
| 57 | *sucAB* | 840555..843356 | 843371..844579 | 15 | 13.33 | 2-oxoglutarate dehydrogenase |
| 58 | *dadAX* | 1894120..1895190 | 1895204..1896502 | 14 | 28.57 | amino acid metabolism |
| 59 | *flgFG* | 1256840..1257595 | 1257609..1258391 | 14 | 28.57 | flagella biosynthesis |
| 60 | *ubiCA* | 4477409..4477906 | 4477920..4478792 | 14 | 35.71 | chorismate pyruvate lyase |
| 61 | *atpHA* | 4099310..4100851 | 4100864..4101397 | 13 | 69.23 | ATP synthase |
| 62 | *eutKR* | 2588826..2589320 | 2589333..2589992 | 13 | 69.23 | ethanolamine utilization protein |
| 63 | *fabDG* | 1274081..1275010 | 1275023..1275757 | 13 | 38.46 | S-malonyltransferase |
| 64 | *fdoGH* | 4266598..4267500 | 4267513..4269927 | 13 | 61.54 | formate dehydrogenase |
| 65 | *manYZ* | 673126..673980 | 673983..674726 | 13 | 53.85 | mannose metabolism |
| 66 | *pyrBI* | 4725486..4725947 | 4725960..4726895 | 13 | 46.15 | aspartate biosynthesis |
| 67 | *rpsC/rplP* | 3615117..3615527 | 3615540..3616241 | 13 | 69.23 | ribosomal units |
| 68 | *acrEF* | 3584243..3585400 | 3585412..3588525 | 12 | 50.00 | acriflavin resistance protein |
| 69 | *cyoDE* | 533857..534747 | 534759..535088 | 12 | 66.67 | cytochrome metabolism |
| 70 | *flgCD* | 1254467..1254871 | 1254883..1255581 | 12 | 66.67 | flagella biosynthesis |
| 71 | *flgHI* | 1258479..1259144 | 1259156..1260253 | 12 | 45.45 | flagella biosynthesis |
| 72 | *oppCD* | 1829139..1830146 | 1830158..1831066 | 12 | 58.33 | oligopeptide transport system |
| 73 | *proBA* | 366286..367389 | 367401..368651 | 12 | 50.00 | gamma-glutamyl kinase |
| 74 | *accBC* | 3573234..3573704 | 3573715..3575064 | 11 | 63.64 | acetyl-CoA carboxyl carrier |
| 75 | *araBA* | 118078..119580 | 119591..121300 | 11 | 54.55 | arabisone operon |
| 76 | *citEF* | 722639..724168 | 724178..725086 | 11 | 54.55 | citrate metabolism |
| 77 | *frdCD* | 4604467..4604826 | 4604837..4605232 | 11 | 72.73 | fumarate reductase |
| 78 | *leuCD* | 129058..129663 | 129674..131074 | 11 | 45.45 | isopropylmalate dehydratase subunit |
| 79 | *nuoIJ* | 2453464..2454018 | 2454029..2454571 | 11 | 63.64 | NADH dehydrogenase |
| 80 | *rplC/rplD* | 3618039..3618644 | 3618655..3619284 | 11 | 54.55 | ribosomal units |
| 81 | *entCE* | 695065..696240 | 696250..697860 | 10 | 50.00 | isochorismate synthase |
| 82 | *fadBA* | 4211011..4212174 | 4212184..4214373 | 10 | 50.00 | fatty acid metabolism |
| 83 | *glyQS* | 3862647..3864716 | 3864726..3865637 | 10 | 80.00 | glycyl-tRNA synthetase |
| 84 | *malPQ* | 3694897..3696975 | 3696985..3699378 | 10 | 80.00 | maltodextrin phosphorylase |
| 85 | *rplF/rplR* | 3611603..3611956 | 3611966..3612499 | 10 | 40.00 | ribosomal units |
| 86 | *trpCB* | 1814226..1815584 | 1815594..1816787 | 10 | 30.00 | trp operon |
| 87 | *glnLG* | 4235612..4237021 | 4237030..4238079 | 9 | 33.33 | nitrogen regulation |
| 88 | *ruvAB* | 1972565..1973575 | 1973584..1974195 | 9 | 44.44 | Holliday junction subunit |
| 89 | *sspAB* | 3533183..3533683 | 3533689..3534327 | 6 | 50.00 | starvation protein |

**Supplementary table 3. Methodology for calculation of base substitutions at intra operon IGRs.**

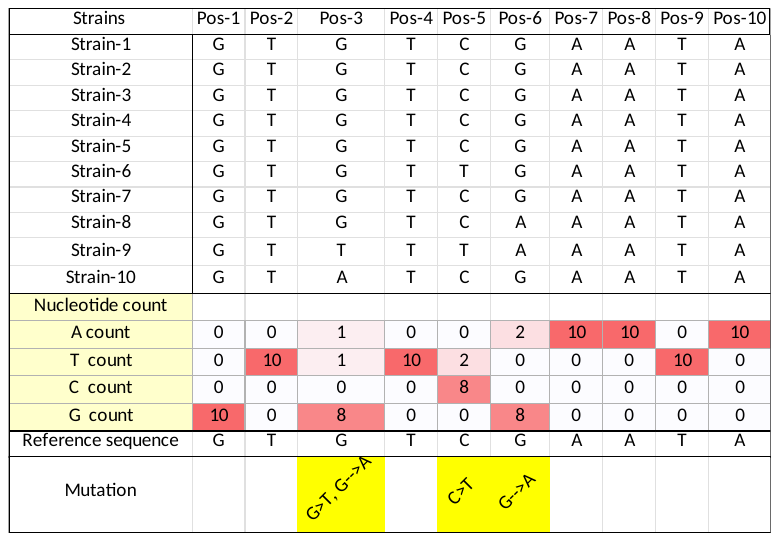

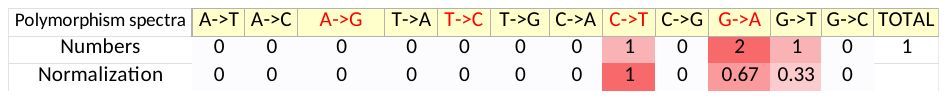

A hypothetical intra operon IGRs having 10bp in size is presented here to show the procedure of reference sequence derivation and calculation of mutations in 10 strains, which was followed in our dataset for the work. The nucleotide count sub-section shows the nucleotide counts of individual columns/positions. At position 1 the most frequent nucleotide was found to be G, hence in reference sequence row, G was mentioned under column/position 1. Similarly, the remaining positions were calculated. At position 3, the frequency of G was found to be 8 and frequency of T and A were found to be 1 each, A and C counts were not found. Hence under position-3, reference sequence row G was mentioned. But the mutation row was mentioned as G🡪T, G🡪A indicating mutations from the most frequent G to the least frequents (T and A) in that position. At position 5, the most frequent nucleotide was found to be C followed by T, having nucleotide counts of 8 and 2 respectively. Similarly at position 6, the most frequent nucleotide was found to be G followed by A. So, position 5 and 6 were indicating C🡪T and G🡪A mutations respectively. Hence it can be said that for this certain scenario we found 4 mutations out of 10 positions. Further this mutation data was considered for the construction of substitution spectra. This methodology was followed for the derivation of reference sequence in our sample intra operon IGRs. Further, in the polymorphism spectra analysis the total numbers of base substitutions can be found along with the normalization value. As described in the Materials and methods section the normalization was done by considering the number of certain base substitution and nucleotide count of the originating nucleotide. Here C🡪T was found to be 1 in number and C nucleotide was found to be 1. Hence the normalization was done by considering the number of C🡪T substitution and dividing it with the number of C. Hence 1/1 =1, that was mentioned in normalization row against C🡪T substitution column. The remaining substitutions were also calculated by this procedure for the entire dataset.

**Supplementary table 4. Collection of intra operon IGRs having base substitutions to find out total spectra in *E. coli*.**

| Sl no. | Operons | A→T | A→C | A→G | T→A | T→C | T→G | C→A | C→T | C→G | G→A | G→T | G→C | Total |
| --- | --- | --- | --- | --- | --- | --- | --- | --- | --- | --- | --- | --- | --- | --- |
| 1 | *tnaAB* | 3 | 1 | 0 | 3 | 1 | 0 | 1 | 0 | 0 | 0 | 0 | 0 | 9 |
|  |  | 0.103 | 0.034 | 0 | 0.120 | 0.040 | 0.000 | 0.045 | 0.000 | 0 | 0 | 0 | 0 |  |
| 2 | *degQS* | 0 | 0 | 0 | 0 | 1 | 1 | 0 | 1 | 0 | 0 | 0 | 0 | 3 |
|  |  | 0 | 0 | 0 | 0.000 | 0.034 | 0.034 | 0 | 0.043 | 0 | 0 | 0 | 0 |  |
| 3 | *dnaKJ* | 2 | 0 | 2 | 1 | 1 | 0 | 0 | 0 | 0 | 0 | 0 | 0 | 6 |
|  |  | 0.08 | 0 | 0.08 | 0.048 | 0.048 | 0 | 0 | 0 | 0 | 0 | 0 | 0 |  |
| 4 | *cadBA* | 0 | 0 | 0 | 0 | 0 | 0 | 0 | 1 | 0 | 0 | 0 | 0 | 1 |
|  |  | 0 | 0 | 0 | 0 | 0 | 0.000 | 0 | 0.056 | 0 | 0 | 0 | 0 |  |
| 5 | *rpoBC* | 0 | 0 | 0 | 0 | 0 | 1 | 0 | 0 | 0 | 0 | 0 | 0 | 1 |
|  |  | 0 | 0 | 0 | 0 | 0 | 0.063 | 0 | 0 | 0 | 0 | 0 | 0 |  |
| 6 | *manXY* | 1 | 0 | 0 | 1 | 0 | 0 | 0 | 1 | 1 | 0 | 0 | 0 | 4 |
|  |  | 0.050 | 0.000 | 0 | 0.048 | 0.000 | 0 | 0 | 0.091 | 0.091 | 0.000 | 0.000 | 0 |  |
| 7 | *lacYA* | 1 | 1 | 0 | 0 | 3 | 0 | 0 | 0 | 1 | 1 | 1 | 0 | 8 |
|  |  | 0.083 | 0.083 | 0 | 0 | 0.158 | 0 | 0 | 0 | 0.067 | 0.056 | 0.056 | 0 |  |
| 8 | *nagBA* | 2 | 0 | 0 | 2 | 0 | 0 | 0 | 2 | 0 | 0 | 0 | 0 | 6 |
|  |  | 0.100 | 0.000 | 0 | 0.167 | 0.000 | 0 | 0 | 0.143 | 0.000 | 0.000 | 0.000 | 0 |  |
| 9 | *nhaAR* | 0 | 0 | 0 | 0 | 0 | 1 | 0 | 2 | 0 | 1 | 0 | 0 | 4 |
|  |  | 0 | 0 | 0 | 0 | 0.000 | 0.071 | 0.000 | 0.154 | 0 | 0.059 | 0 | 0 |  |
| 10 | *creCD* | 0 | 0 | 0 | 1 | 0 | 0 | 0 | 2 | 0 | 0 | 0 | 0 | 3 |
|  |  | 0 | 0 | 0 | 0.067 | 0 | 0.000 | 0 | 0.118 | 0 | 0 | 0 | 0 |  |
| 11 | *deoBD* | 0 | 0 | 0 | 1 | 0 | 1 | 0 | 1 | 0 | 2 | 0 | 0 | 5 |
|  |  | 0 | 0 | 0 | 0.067 | 0.000 | 0.067 | 0.000 | 0.1 | 0 | 0.133 | 0 | 0 |  |
| 12 | *lacZY* | 0 | 0 | 0 | 0 | 1 | 1 | 1 | 0 | 0 | 2 | 0 | 0 | 5 |
|  |  | 0 | 0 | 0 | 0 | 0.071 | 0.071 | 0.083 | 0 | 0 | 0.154 | 0 | 0 |  |
| 13 | *atpAG* | 0 | 0 | 0 | 1 | 0 | 0 | 0 | 0 | 0 | 0 | 0 | 0 | 1 |
|  |  | 0 | 0 | 0.000 | 0.111 | 0.000 | 0.000 | 0.000 | 0.000 | 0 | 0.000 | 0.000 | 0 |  |
| 14 | *deoAB* | 0 | 0 | 0 | 0 | 1 | 0 | 0 | 0 | 0 | 0 | 0 | 0 | 1 |
|  |  | 0 | 0 | 0.000 | 0.000 | 0.077 | 0.000 | 0.000 | 0.000 | 0 | 0.000 | 0.000 | 0 |  |
| 15 | *fhuAC* | 0 | 0 | 1 | 1 | 2 | 0 | 0 | 3 | 0 | 0 | 0 | 0 | 7 |
|  |  | 0 | 0 | 0.167 | 0.053 | 0.105 | 0.000 | 0.000 | 0.250 | 0 | 0.000 | 0.000 | 0 |  |
| 16 | *livKH* | 1 | 0 | 0 | 0 | 1 | 0 | 0 | 1 | 0 | 4 | 1 | 0 | 8 |
|  |  | 0.1 | 0 | 0.000 | 0 | 0.091 | 0.000 | 0.000 | 0.083 | 0 | 0.308 | 0.077 | 0 |  |
| 17 | *hicAB* | 0 | 0 | 2 | 0 | 1 | 0 | 0 | 0 | 0 | 0 | 0 | 0 | 3 |
|  |  | 0 | 0 | 0.105 | 0 | 0.100 | 0.000 | 0.000 | 0 | 0 | 0.000 | 0 | 0 |  |
| 18 | *fliAZ* | 0 | 0 | 0 | 0 | 0 | 0 | 0 | 0 | 0 | 1 | 0 | 0 | 1 |
|  |  | 0 | 0 | 0 | 0 | 0.000 | 0.000 | 0.000 | 0 | 0 | 0.091 | 0 | 0 |  |
| 19 | *nrfAB* | 0 | 0 | 0 | 0 | 1 | 0 | 1 | 0 | 0 | 1 | 0 | 0 | 3 |
|  |  | 0 | 0 | 0 | 0 | 0.167 | 0.000 | 0.077 | 0 | 0 | 0.077 | 0 | 0 |  |
| 20 | *groSL* | 0 | 0 | 0 | 0 | 0 | 0 | 0 | 0 | 0 | 0 | 1 | 0 | 1 |
|  |  | 0 | 0 | 0 | 0 | 0.000 | 0.000 | 0.000 | 0 | 0 | 0.000 | 0.125 | 0 |  |
| 21 | *ptsHI* | 0 | 0 | 0 | 0 | 1 | 0 | 1 | 0 | 0 | 0 | 0 | 0 | 2 |
|  |  | 0 | 0 | 0.000 | 0.000 | 0.071 | 0 | 0.143 | 0.000 | 0.000 | 0.000 | 0.000 | 0 |  |
| 22 | *xapAB* | 0 | 0 | 1 | 2 | 0 | 0 | 1 | 2 | 0 | 1 | 0 | 0 | 7 |
|  |  | 0 | 0 | 0.143 | 0.083 | 0.000 | 0 | 0.063 | 0.125 | 0.000 | 0.091 | 0.000 | 0 |  |
| 23 | *nuoFG* | 0 | 0 | 0 | 2 | 1 | 0 | 0 | 0 | 0 | 0 | 0 | 0 | 3 |
|  |  | 0 | 0 | 0.000 | 0.154 | 0.063 | 0 | 0.000 | 0.000 | 0.000 | 0.000 | 0.000 | 0 |  |
| 24 | *csgBA* | 0 | 0 | 0 | 1 | 0 | 0 | 1 | 1 | 0 | 0 | 0 | 0 | 3 |
|  |  | 0 | 0 | 0.000 | 0.063 | 0.000 | 0 | 0.143 | 0.143 | 0.000 | 0.000 | 0.000 | 0 |  |
| 25 | *prpDE* | 0 | 0 | 0 | 0 | 1 | 0 | 0 | 1 | 0 | 1 | 1 | 0 | 4 |
|  |  | 0 | 0 | 0.000 | 0 | 0.167 | 0 | 0 | 0.167 | 0.000 | 0.077 | 0.077 | 0 |  |
| 26 | *agaBC* | 0 | 0 | 1 | 0 | 0 | 0 | 1 | 0 | 0 | 0 | 0 | 0 | 2 |
|  |  | 0 | 0 | 0.071 | 0 | 0.000 | 0 | 0.25 | 0.000 | 0.000 | 0 | 0 | 0 |  |
| 27 | *prpCD* | 0 | 0 | 0 | 0 | 0 | 1 | 0 | 0 | 0 | 1 | 0 | 0 | 2 |
|  |  | 0.000 | 0 | 0 | 0 | 0 | 0.2 | 0 | 0 | 0 | 0.25 | 0.000 | 0 |  |
| 28 | *tdcDE* | 0 | 0 | 1 | 0 | 1 | 0 | 0 | 0 | 0 | 0 | 0 | 0 | 2 |
|  |  | 0.000 | 0 | 0.066 | 0 | 0.111 | 0 | 0 | 0 | 0 | 0 | 0.000 | 0 |  |
| 29 | *fucPI* | 0 | 0 | 0 | 1 | 0 | 0 | 0 | 0 | 0 | 0 | 0 | 0 | 1 |
|  |  | 0.000 | 0 | 0 | 0.1 | 0 | 0 | 0 | 0 | 0 | 0 | 0.000 | 0 |  |
| 30 | *caiTA* | 0 | 0 | 0 | 0 | 0 | 1 | 0 | 0 | 0 | 0 | 2 | 0 | 3 |
|  |  | 0.000 | 0 | 0 | 0 | 0 | 0.071 | 0 | 0 | 0 | 0 | 0.333 | 0 |  |
| 31 | *marAB* | 1 | 0 | 0 | 0 | 0 | 0 | 0 | 0 | 0 | 0 | 1 | 0 | 2 |
|  |  | 0.091 | 0 | 0 | 0 | 0 | 0 | 0 | 0 | 0 | 0 | 0.143 | 0 |  |
| 32 | *atoEB* | 0 | 0 | 1 | 1 | 0 | 0 | 1 | 2 | 0 | 0 | 0 | 0 | 5 |
|  |  | 0 | 0 | 0.077 | 0.333 | 0 | 0 | 0.125 | 0.25 | 0 | 0 | 0 | 0 |  |
| 33 | *lsrBF* | 0 | 1 | 0 | 0 | 0 | 0 | 0 | 0 | 0 | 1 | 0 | 0 | 2 |
|  |  | 0 | 0.125 | 0 | 0 | 0 | 0 | 0 | 0 | 0 | 0.167 | 0 | 0 |  |
| 34 | *cmtBA* | 0 | 1 | 1 | 0 | 0 | 0 | 0 | 0 | 0 | 1 | 0 | 0 | 3 |
|  |  | 0 | 0.1 | 0.1 | 0 | 0 | 0 | 0 | 0 | 0 | 0.143 | 0 | 0 |  |
| 35 | *marRA* | 0 | 0 | 0 | 0 | 0 | 0 | 0 | 1 | 0 | 0 | 0 | 0 | 1 |
|  |  | 0 | 0 | 0 | 0 | 0 | 0 | 0 | 0.5 | 0 | 0 | 0 | 0 |  |
| 36 | *eutBC* | 0 | 0 | 0 | 0 | 0 | 0 | 0 | 0 | 0 | 1 | 0 | 0 | 1 |
|  |  | 0 | 0 | 0 | 0 | 0 | 0 | 0 | 0 | 0 | 0.2 | 0 | 0 |  |
| 37 | *atpDC* | 0 | 0 | 0 | 0 | 0 | 0 | 1 | 0 | 0 | 0 | 0 | 0 | 1 |
|  |  | 0 | 0 | 0 | 0 | 0 | 0 | 0.142 | 0 | 0 | 0 | 0 | 0 |  |
| 38 | *fliIJ* | 0 | 0 | 0 | 0 | 0 | 0 | 0 | 1 | 0 | 0 | 0 | 0 | 1 |
|  |  | 0 | 0 | 0 | 0 | 0 | 0 | 0 | 0.2 | 0 | 0 | 0 | 0 |  |
| 39 | *cusFB* | 0 | 0 | 1 | 0 | 0 | 0 | 0 | 1 | 0 | 0 | 1 | 0 | 3 |
|  |  | 0 | 0 | 0.2 | 0 | 0 | 0 | 0 | 0.333 | 0 | 0 | 0.25 | 0 |  |
| 40 | *mglAC* | 0 | 0 | 0 | 0 | 0 | 0 | 0 | 1 | 0 | 0 | 0 | 0 | 1 |
|  |  | 0 | 0 | 0 | 0 | 0 | 0 | 0 | 0.2 | 0 | 0 | 0 | 0 |  |
| 41 | *hdeAB* | 0 | 1 | 0 | 0 | 1 | 0 | 1 | 0 | 0 | 2 | 1 | 0 | 6 |
|  |  | 0 | 0.027 | 0 | 0 | 0.036 | 0 | 0.063 | 0 | 0 | 0.087 | 0.043 | 0 |  |
| 42 | *araGH* | 1 | 0 | 0 | 0 | 0 | 0 | 0 | 0 | 0 | 0 | 0 | 0 | 1 |
|  |  | 0.333 | 0 | 0 | 0 | 0 | 0 | 0 | 0 | 0 | 0 | 0 | 0 |  |
| 43 | *pyrBI* | 0 | 0 | 0 | 0 | 1 | 0 | 0 | 0 | 0 | 0 | 0 | 0 | 1 |
|  |  | 0 | 0 | 0 | 0 | 0.166 | 0 | 0 | 0 | 0 | 0 | 0 | 0 |  |
| 44 | *tauAB* | 0 | 0 | 1 | 0 | 0 | 0 | 0 | 0 | 0 | 0 | 0 | 0 | 1 |
|  |  | 0 | 0 | 0.2 | 0 | 0 | 0 | 0 | 0 | 0 | 0 | 0 | 0 |  |
| 45 | *gabDT* | 0 | 0 | 0 | 0 | 1 | 0 | 0 | 0 | 0 | 0 | 0 | 0 | 1 |
|  |  | 0 | 0 | 0 | 0 | 0.5 | 0 | 0 | 0 | 0 | 0 | 0 | 0 |  |
| 46 | *cusBA* | 0 | 0 | 1 | 0 | 0 | 0 | 0 | 0 | 0 | 0 | 0 | 0 | 1 |
|  |  | 0 | 0 | 0.166 | 0 | 0 | 0 | 0 | 0 | 0 | 0 | 0 | 0 |  |
| 47 | *fabDG* | 0 | 1 | 0 | 0 | 0 | 0 | 0 | 0 | 0 | 0 | 0 | 0 | 1 |
|  |  | 0 | 0.142 | 0 | 0 | 0 | 0 | 0 | 0 | 0 | 0 | 0 | 0 |  |
| 48 | *manYZ* | 0 | 0 | 0 | 0 | 0 | 0 | 0 | 1 | 0 | 0 | 0 | 0 | 1 |
|  |  | 0 | 0 | 0 | 0 | 0 | 0 | 0 | 0.5 | 0 | 0 | 0 | 0 |  |
| 49 | *araBA* | 0 | 0 | 1 | 0 | 0 | 0 | 0 | 0 | 0 | 0 | 0 | 0 | 1 |
|  |  | 0 | 0 | 1 | 0 | 0 | 0 | 0 | 0 | 0 | 0 | 0 | 0 |  |
| 50 | *proBA* | 1 | 0 | 0 | 0 | 1 | 0 | 0 | 0 | 0 | 0 | 0 | 0 | 2 |
|  |  | 0.5 | 0 | 0 | 0 | 1 | 0 | 0 | 0 | 0 | 0 | 0 | 0 |  |
| 51 | *flgCD* | 0 | 0 | 0 | 0 | 0 | 0 | 0 | 0 | 0 | 1 | 0 | 0 | 1 |
|  |  | 0 | 0 | 0 | 0 | 0 | 0 | 0 | 0 | 0 | 0.25 | 0 | 0 |  |
| 52 | *acrEF* | 0 | 0 | 0 | 0 | 1 | 0 | 0 | 0 | 0 | 0 | 0 | 0 | 1 |
|  |  | 0 | 0 | 0 | 0 | 0.33 | 0 | 0 | 0 | 0 | 0 | 0 | 0 |  |
| 53 | *malPQ* | 0 | 0 | 0 | 0 | 0 | 0 | 0 | 0 | 0 | 1 | 0 | 0 | 1 |
|  |  | 0 | 0 | 0 | 0 | 0 | 0 | 0 | 0 | 0 | 0.5 | 0 | 0 |  |
| 54 | *codBA* | 0 | 0 | 0 | 0 | 0 | 0 | 0 | 0 | 0 | 1 | 0 | 0 | 1 |
|  |  | 0 | 0 | 0 | 0 | 0 | 0 | 0 | 0 | 0 | 0.33 | 0 | 0 |  |
| 55 | *leuCD* | 0 | 0 | 0 | 0 | 0 | 0 | 0 | 0 | 0 | 0 | 0 | 1 | 1 |
|  |  | 0 | 0 | 0 | 0 | 0 | 0 | 0 | 0 | 0 | 0 | 0 | 0.25 |  |
| 56 | *flgHI* | 0 | 0 | 0 | 0 | 0 | 0 | 0 | 0 | 0 | 1 | 0 | 0 | 1 |
|  |  | 0 | 0 | 0 | 0 | 0 | 0 | 0 | 0 | 0 | 0.25 | 0 | 0 |  |
| 57 | *entCE* | 0 | 0 | 0 | 0 | 0 | 0 | 0 | 1 | 0 | 0 | 0 | 0 | 1 |
|  |  | 0 | 0 | 0 | 0 | 0 | 0 | 0 | 1 | 0 | 0 | 0 | 0 |  |
|  | ***Total spectra*** |  |  |  |  |  |  |  |  |  |  |  |  |  |
| **Number** |  | **15** | **5** | **20** | **16** | **17** | **7** | **11** | **31** | **1** | **19** | **8** | **2** | **152** |
| **Normalization** |  | **0.014** | **0.005** | **0.019** | **0.020** | **0.021** | **0.009** | **0.017** | **0.049** | **0.002** | **0.021** | **0.009** | **0.002** |  |

In case of *E. coli* out of 134 intra operon IGRs, we found substitutions in 57 IGRs. Finally, we considered the 57 IGRs for final spectra constructions. It was also considered for the *S. enterica* species.

**Supplementary table 5. The total numbers of individual base substitution present in inter operon IGRs and intra operon IGRs present in *E. coli* and *S. enterica* are listed.**

| Substitutions | *Numbers of individual base substitutions* | | | |
| --- | --- | --- | --- | --- |
|  | Inter operon IGRs | | Intra operon IGRs | |
|  | *E. coli* | *S. enterica* | *E. coli* | *S. enterica* |
| A->T | 1732 | 1932 | 15 | 2 |
| A->C | 1185 | 1827 | 5 | 0 |
| A->G | 3573 | 6348 | 20 | 20 |
| T->A | 1759 | 1853 | 16 | 10 |
| T->C | 3642 | 6359 | 17 | 10 |
| T->G | 1154 | 1805 | 7 | 3 |
| C->A | 1742 | 3398 | 11 | 7 |
| C->T | 5107 | 11117 | 31 | 56 |
| C->G | 666 | 1106 | 1 | 1 |
| G->A | 5103 | 11037 | 19 | 28 |
| G->T | 1726 | 3354 | 8 | 16 |
| G->C | 667 | 1060 | 2 | 2 |
| Total | 28056 | 51196 | 152 | 155 |

1. Start and end coordinates and function and details of intra operon IGRs were extracted based on data availability in National Center of Biotechnology Information (NCBI) by considering the reference strain ***Escherichia coli* str. K12 substr. MG1655 (**NCBI: txid511145)
2. Start and end coordinates and function and details of intra operon IGRs were extracted based on data availability in National Center of Biotechnology Information (NCBI) by considering the reference strain ***Salmonella enterica subsp. enterica serovar Typhimurium str. D23580* (**NCBI: txid568708)
